# An unbroken network of interactions connecting flagellin domains is required for motility in viscous environments

**DOI:** 10.1101/2022.11.07.515560

**Authors:** Marko Nedeljković, Sandra Postel, Daniel Bonsor, Yingying Xing, Neil Jacob, William J. Schuler, Eric J. Sundberg

## Abstract

In its simplest form, bacterial flagellar filaments are composed of flagellin proteins with just two helical inner domains, which together comprise the filament core. Although this minimal filament is sufficient to provide motility in many flagellated bacteria, most bacteria produce flagella composed of flagellin proteins with one or more outer domains arranged in a variety of supramolecular architectures radiating from the inner core. Flagellin outer domains are known to be involved in adhesion, proteolysis and immune evasion but have not been thought to be required for motility. Here we show that in the *Pseudomonas aeruginosa* POA1 strain, a bacterium that forms a ridged filament on account of the arrangement of the two outer domains of its flagellin protein, motility is categorically dependent on these flagellin outer domains. Moreover, a comprehensive network of intermolecular interactions connecting the inner domains to the outer domains, the outer domains to one another, and the outer domains back to the inner domain filament core, is required for motility. This inter-domain connectivity confers PAO1 flagella with increased stability, essential for its motility in viscous environments. Additionally, we find that such ridged flagellar filaments are not unique to *Pseudomonas* but are, instead, present throughout diverse bacterial phyla.

## INTRODUCTION

The bacterial flagellum is a complex and dynamic nanomachine that propels bacteria through liquids. In pathogenic species, motility provided by flagella is critical for host colonization and infection, and flagellar filaments are recognized as important virulence factors involved in adherence, toxin delivery, biofilm formation and activation of innate immunity [1–8]. The number of bacterial flagella attached to the cell body varies among species, from one, as in *Pseudomonas aeruginosa*, to many, as in *Escherichia coli* and *Salmonella enterica* serovar Typhimurium [9]. Each flagellum consists of a membrane-embedded basal body that functions as a motor, as well as a hook and a long filament [10, 11]. Torque generated by the basal body is transferred by the hook to the filament, rotation of which provides a thrust that propels bacteria through liquid.

The flagellar filament, a tubular structure composed of thousands of copies of the protein flagellin (FliC in many bacteria), exhibits helical symmetry [12]. One FliC molecule is copied 11 times along the screw axis, making two turns, before it reaches the position directly above this starting position. Such an arrangement results in stacks of FliC along the filament axis called protofilaments, functional units of the filament that actively switch between left-(L) and right-(R) handed forms independently of their neighboring protofilament. Different combinations of protofilaments determine the twist of the filament and, consequently, swimming speed [13, 14]. If all protofilaments are in the same form, they produce a straight filament that does not create thrust, resulting in immotile bacteria.

The filament is capped at its distal end by an oligomeric stool-like structure comprised of five or six copies of the protein FliD [15, 16]. FliC molecules are synthesized in the cytoplasm and exported through the flagellar type III secretion system at the base of the flagellum [17]. Since the filament extends from its distal end, FliC must be transported through the central channel of the filament, which measures approximately 25 Å in diameter. Thus, FliC must be at least partially unfolded while passing through the channel and, once it reaches the FliD cap, it attains the correct structure and is positioned at the tip of the filament [17, 18]. This process is still not fully elucidated, but involves direct interactions between incoming FliC subunits and the oligomeric FliD cap in a species-specific manner [19].

The flagellar system of *Salmonella* has been studied as the archetypal model of bacterial flagella for decades. The first high-resolution flagellin protein structure was from that of *S.* Typhimurium [20], revealing a boomerang-shaped molecule with four domains – the D0 and D1 inner domains that are predominantly α-helical and comprise the filament core, and the propeller-shaped D2 and D3 outer domains that protrude from the filament at an angle of approximately 90 degrees to the filament axis [20, 21]. In *S.* Typhimurium flagellar filaments, the outer domains are splayed outward from the filament core such that no extensive interactions are made between the D2 and D3 domains of neighboring subunits along the protofilament [20, 22]. This is corroborated by mutational studies that have shown that large truncations of the *S.* Typhimurium outer domains are tolerated without abolishing motility [23]. Recently, structures of flagellins and filaments from other species have become available, confirming that the inner core, composed of D0 and D1 domains, is structurally conserved, while there is extensive variability in the sequence, length, structure and arrangement of outer domains in FliC [24–26]. In some motile bacteria, the outer domains are completely absent, as observed for *Bacillus subtilis* and *Kurthia* sp. [27, 28]. A recent series of high-resolution structures of so-called complex filaments showed that outer domains can oligomerize to generate helical threads or mesh-like sheath structures that serve to stabilize the filament form [26]. Thus, flagellated motile bacteria all share a highly conserved inner core decorated by a widely divergent collection of outer domains.

Although the mechanism for switching between L- and R-handed conformations of the filament is not fully understood, the inner core has been shown to be responsible for this change [27]. Mutations that lock the *Salmonella* filament in L or R form are located in the D1 domain [29]. Equivalent mutations produce the same effect in other species, including *B. subtilis*, *P. aeruginosa* and *Campylobacter jejuni* [25, 27]. Outer domains, on the other hand, have not been shown to contribute to hand-switching or motility and, instead, are thought to mediate many of the other roles in which flagella are implicated, such as adherence and biofilm formation. However, the growing catalog of diverse flagellar filament supramolecular architectures suggests that FliC outer domains could, in certain bacteria, play a critical role in motility. The previously reported 4.2 Å cryo-EM structures of L- and R-handed filaments from the *P. aeruginosa* strain PAO1 revealed that the overall shape of this filament was ridged, unlike the splayed geometry of the *S.* Typhimurium filament [27]. In the former, the D2 and D3 outer domains of FliC adopt an end-on-end compact fold along the filament axis, resulting in ridges along the filament where they are present and clefts along the filament where they are absent. Due to low local resolution (>10 Å) of the D2 and D3 domains in the cryo-EM structure, the intermolecular interactions maintaining the ridges, as well as connecting the ridges to the inner core, were not resolved. In order to obtain a complete model of the PAO1 filament and to study the functional consequences of the “ridged” structural organization, we turned to X-ray crystallography to resolve the structure of the missing D2-D3 domains and, subsequently, performed a whole-atom reconstruction of the *P. aeruginosa* PAO1 filament. Based on the resulting model, we identified three interfaces that the D2 and D3 outer domains form with one another as well as with the D1 inner domain. Combining genetic tools, electron microscopy and swimming motility assays, we showed that these interfaces form a connected network of inter-domain interfaces that are critical not only for the structural integrity of the filament, but also for motility of the bacterium. Using bioinformatic tools, we found that ridged flagellar filaments similar to that in *P. aeruginosa* PAO1 are common throughout the bacterial kingdom and that such interconnected filament architectures may provide an advantage to bacteria in generating the thrust needed to swim through liquids of high viscosity.

## RESULTS

### High-resolution structure of the *P. aeruginosa* PAO1 FliC D2 and D3 domains

Although the cryo-EM structure of the PAO1 filament achieved near-atomic resolution for the inner core of the filament, the resolution of the outer domains was substantially lower, prohibiting even the tracing of the main chain of the D2 and D3 domains [27]. Thus, we expressed and purified the fragment of FliC between the residues 178-395 that forms the D2 and D3 domains and crystallized both the native protein and its selenomethionine derivative in the P2 space group with 2 molecules in the asymmetric unit. Employing multi-wavelength anomalous dispersion (MAD) phasing methods, we arrived at a 1.47 Å resolution structure revealing the D2 and D3 domains as a contiguous barrel-like structure composed predominantly of β-strands, with D3 positioned directly above D2 along the filament axis towards the distal end of the filament (**Figure 1A, S1A and B, Supplementary table 1**). While D3 in *S.* Typhimurium is inserted into D2 and they form two separated domains, a different topology of the secondary structural elements in PAO1 FliC results in the D2 and D3 domains folding into a single bilobal structure with two distinct surfaces, the concave surface facing the filament core, and the convex surface exposed to the outside. Starting from the N-terminus, the first β-strand is incorporated into the D3 moiety, before the backbone turns downward, forming one half of D2. The sequence between 243 and 353 is entirely a part of D3, after which the polypeptide chain returns to complete the fold of D2. On the concave side of the D2/D3 domain, residues between 233 and 242 belong to the β4-strand that occupies central position spanning both D2 and D3 and together with strands 7, 8, 10 and 11 of D3 and strand 13 of D2 they form an antiparallel β-sheet. The consequence of this organization is the tight packing of D2 and D3 which are oriented along the filament, instead of projecting away from it as in the *S.* Typhimurium.

**Figure 1.**
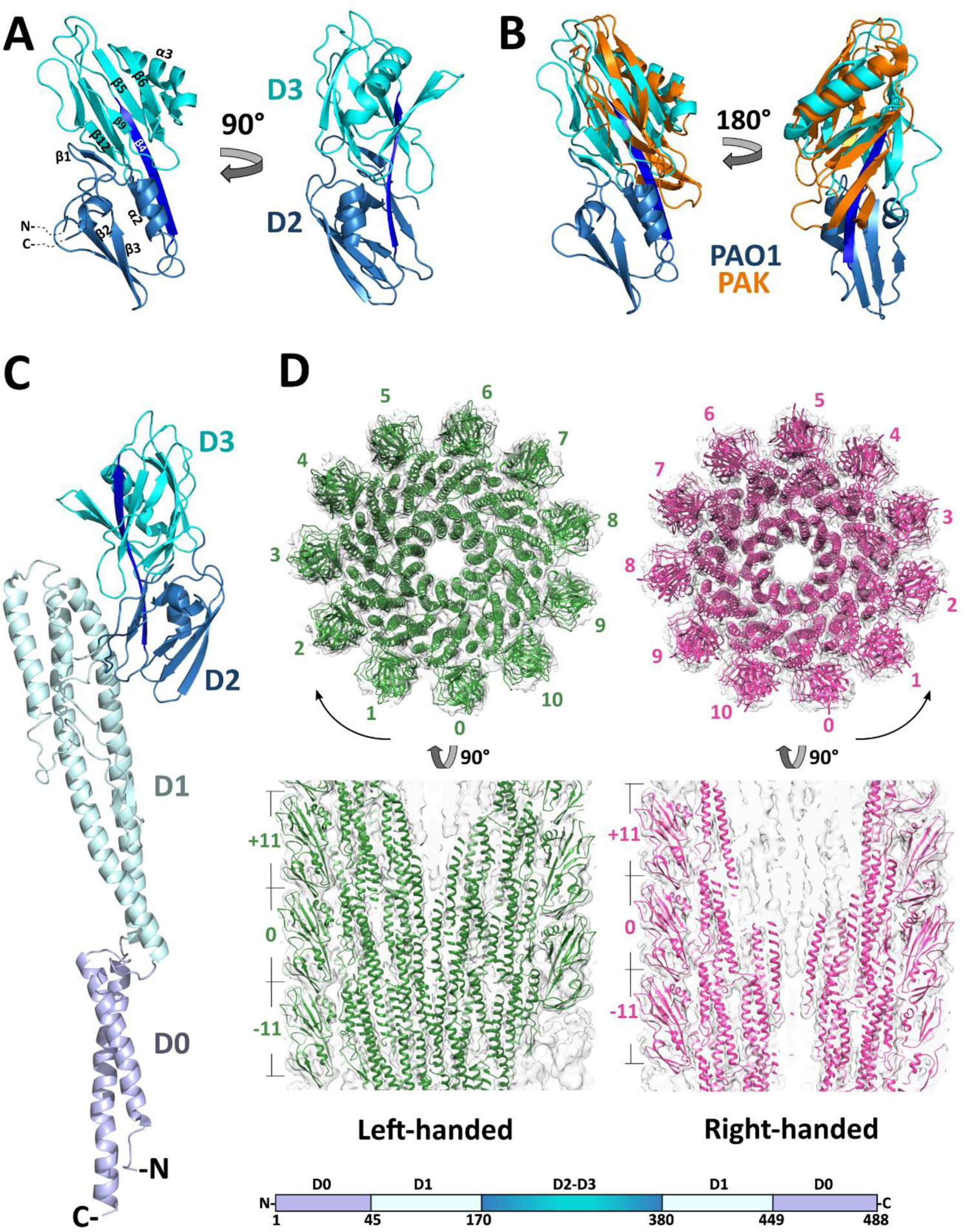
**A** – Crystal structure of the FliC D2/D3 domains from *P. aeruginosa*. Marine blue – D2, cyan – D3, dark blue – common beta-strand. Alpha-helices and beta-strands are numbered in order of appearance in D2/D3 domains, not in full-length FliC; **B** – Superposition of outer domain structures of *P. aeruginosa* PAO1 (color code as in A) and PAK (orange) strains.; **C** – Full-length FliC of PAO1 with schematic representation of FliC protein organization. Colors on the bar correspond to domain colors on the structure; **D** – Cross-sections of the left-handed (green) and right-handed (magenta) filaments of PAO1. Numbers denote the position of FliC subunits in the filament in relation to the reference subunit of 0. Subunits +11 and −11 are directly above and below subunit 0 in the same protofilament.

Two types of FliC proteins are found in *P. aeruginosa*, type A as observed in the PAK strain or type B as seen in *P. aeruginosa* PAO1 [30]. The inner domains display high sequence identity, while the outer domains share poor sequence conservation and size. Compared to the previously determined structure of the type A PAK FliC [24], our type B PAO1 FliC outer domains are oriented similarly relative to the inner core. However, the type A D2 domain of PAK is 90 residues smaller and structurally corresponds to the D3 moiety in PAO1 type B FliC (**Figure 1B**).

### Whole-atom reconstruction of the L- and R-handed filaments in *P. aeruginosa* PAO1

Using the previously determined cryo-EM structure of the L- and R-PAO1 filament, we successfully placed the X-ray crystal structure of the D2 and D3 PAO1 FliC into the residual density and modelled the loops connecting D2-D3 to D1 (**Figure 1C and S1C**). We reconstructed complete filaments using Phenix [31] by applying helical symmetry parameters obtained from the cryo-EM map and subsequently refined this reconstruction using the real-space refinement function in Phenix (**Figure 1D and Supplementary table 2**). Electron density maps for L- and R-handed filaments showed that a protein-protein interface is formed between the D2 and D3 domains of FliC monomers stacked on top of one another in the filament.

As shown in **Figure 2**, these intermolecular D2-D3 interfaces (e.g., between the D3 domain of one FliC subunit, D3^0^, and the D2 domain of the next FliC subunit along the protofilament in the direction of the distal end of the filament, D2^+11^) are similar in size between distinctly handed filaments (331 Å^2^ in L- and 381 Å^2^ in R-handed filament) and consist of residues in helix α2, loop regions 249-253 and 266-269, as well as residue T315. In our models, the glycine turn between strands β2 and β3 of D2^+11^ is positioned directly above helix α2 of D3^0^, engaging in polar contacts with Q277 and S280 in L- and R-filaments, respectively. An additional hydrogen bond between A266 and V207 is present in the L-filament. Apart from the direct contacts within the outer domains, D3 also forms an interface with the β hairpin on D1 domain of the neighboring FliC subunit (D3^0^-D1^+11^ interface) of comparable buried surface area. We identified 4 possible polar contacts between D3^0^ and D1^+11^ in L-handed and none in R-handed conformation. Finally, D2 also forms polar contacts with its own D1 domain (D2^0^-D1^0^ interface). Thus, the ridged flagellar filament of *P. aeruginosa* PAO1 results in three distinct interfaces, moving through FliC subunits and along the protofilament towards the distal end of the filament: one that is intramolecular (D2^0^-D1^0^) and two that are intermolecular (D3^0^-D2^+11^ and D3^0^-D1^+11^).

**Figure 2.**
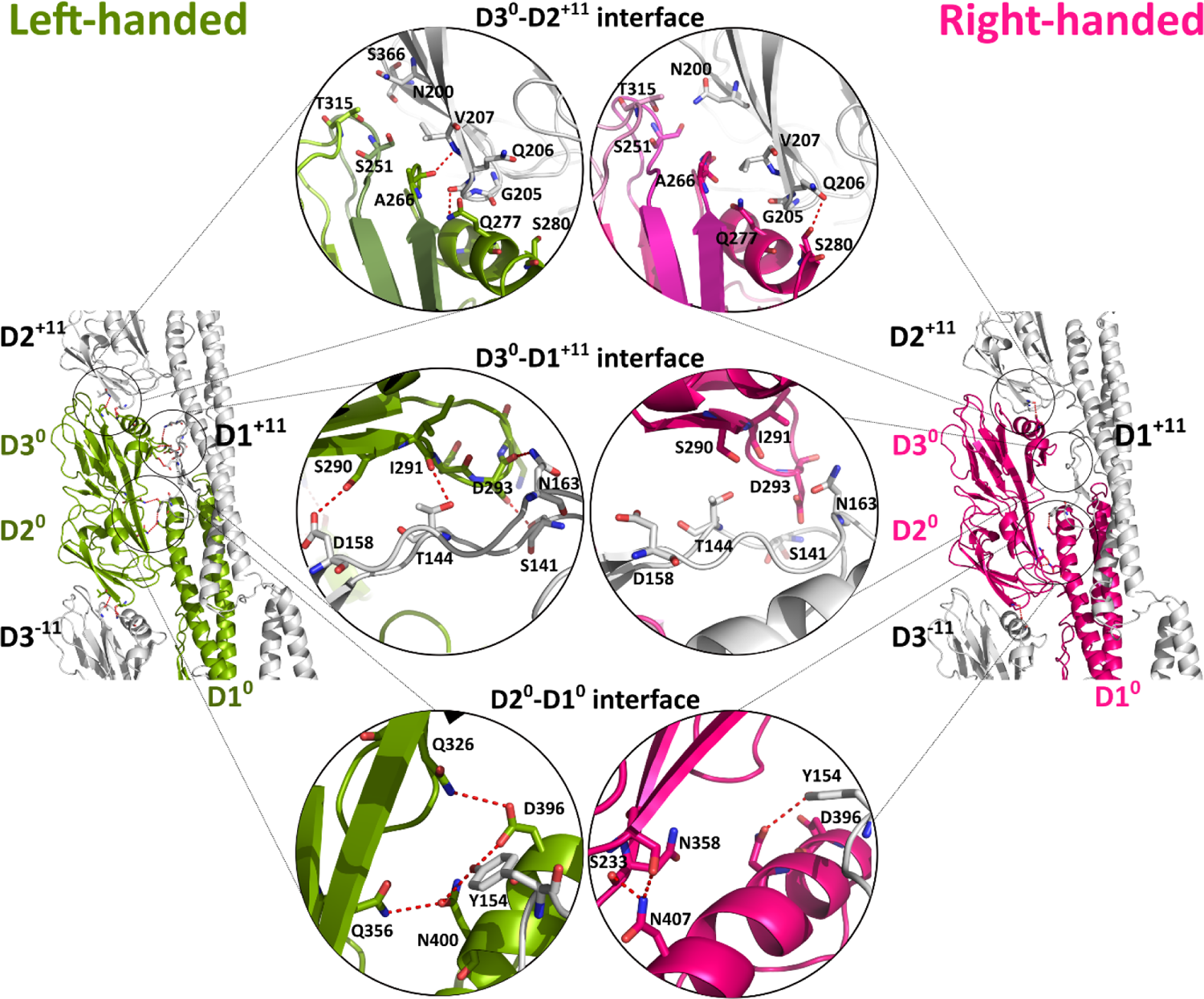
FliC monomer in left-handed (green) and right-handed (magenta) protofilament conformation. Three interfaces that D2^0^ and D3^0^ form with surrounding domains are zoomed in on (highlighted with circles): D3^0^-D2^+11^ (equivalent to D3^-11^-D2^0^), D3^0^-D1^+11^ and D2^0^-D1^0^. Dashed red lines represent hydrogen bonds.

### A network of inter-domain interactions in *P. aeruginosa* PAO1 flagella is required for motility

In order to determine the functional importance of the observed D2/D3 interfaces in the filament, we generated a series of *fli*C mutant genes on a plasmid that we used for *P. aeruginosa* PAO1-*Δfli*C strain complementation. We then observed the effects of mutations on filament formation and tested swimming motility in an agar-based swimming assay by comparing their motile spread to that of the *P. aeruginosa* PAO1-*Δfli*C strain complemented with the wild type gene (**Figures 3, S2A-R and S3**). FliC mutations were of two kinds – in the case of polar interactions involving side chains we mutated residues to alanine, while in the cases of hydrogen bonds involving main chain atoms, as well as loops that were part of the buried surface areas of the inter-domain interfaces, but did not form hydrogen bonds, we deleted a variable number of residues. Our mutational studies indicated that each one of the inter-domain interfaces within *P. aeruginosa* PAO1 filaments – the D2^0^-D1^0^, D3^0^-D2^+11^ and D3^0^-D1^+11^ interfaces – are critical for motility and that, collectively, they form a network of interactions bridging the outer domains to the inner core of the filament that is required for motility, as follows.

**Figure 3.**
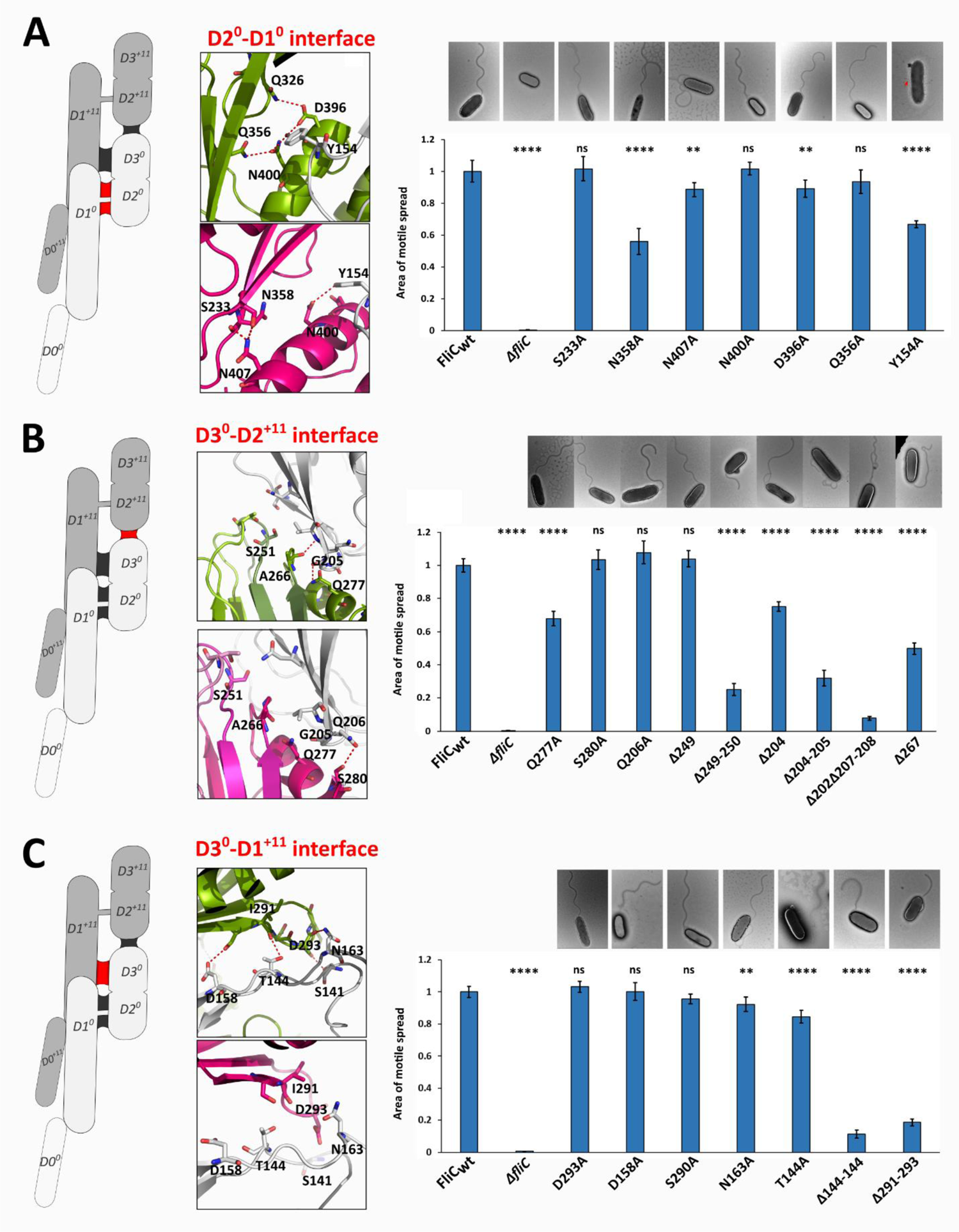
Effects of the FliC mutations on *P. aeruginosa* PAO1 swimming motility and filament formation. Swimming motility analysis and negative-stain EM images of PAO1-*ΔfliC* strain complemented with FliC bearing mutations in the D2^0^-D1^0^ interface (**A**), D3^0^-D2^+11^ interface (**B**), and D3^0^-D1^+11^ interface (**C**). Positions of the interfaces are labeled red on the FliC dimer cartoon on the left. Area of motile spread for each strain is normalized to that of the full-length wild type complemented strain (FliC_WT_). (ns – not significant; ** p < 0.01; **** p < 0.0001).

In the D2^0^-D1^0^ interface, we mutated seven residues that engage in hydrogen bonds and reside on secondary structural elements in our model into alanine. Although also a part of the D1^0^-D1^+11^ interface, Y154 of FliC^+11^ participates in the hydrogen bond network formed by Q326, Q356, D396 and N400, which are located in the D2^0^-D1^0^ interface, and we mutated this residue as well. As observed by electron microscopy, all of these mutations resulted in flagellar filament formation (**Figure 3A**). The majority of these mutations statistically significantly impaired swimming motility compared to wild type. FliC mutants Y154A and N358A exhibited the most substantial decreases in motility, with reductions in motile spread of 35% and 45%, respectively.

In the D3^0^-D2^+11^ interface (**Figure 3B**), single site alanine mutations had either modest or no effect on motility, however, alanine mutation of Q277, which forms a hydrogen bond with G205^+11^ in the L-handed filament, decreased the motile spread by one-third of that of the wild type FliC. Conversely, shortening of the loops by truncation led to significant decreases in motile spread in all but one case. As for the D2^0^-D1^0^ interface, all FliC modifications in the D3^0^-D2^+11^ interface resulted in the formation of flagellar filaments, as observed by electron microscopy. Even in the most extreme case of Δ202Δ207-208, which was nearly entirely immotile, filaments formed. However, the morphology of the filaments for some mutants differed compared to the wild type. While the filaments in Δ249-250 were shorter than the wild type, those of Δ204-205, Δ202Δ207-208 and Δ267 were primarily characterized by the loss of the wavy form observed for wild type filaments, potentially implicating the area around the glycine turn in handedness and/or hand-switching in these filaments. It should be noted that any filament visible in these micrographs, however short they appear, are composed of at least thousands of subunits of FliC extending well beyond the flagellar hook.

While mutations of individual residues in the D3^0^-D1^+11^ interface did not markedly affect PAO1 motility, we observed a profound effect on swimming motility, when either of the two loop regions on D3^0^ or D1^+11^ that engage in hydrogen bonds were truncated in order to abrogate these hydrogen bonding networks (Δ291-293 on D3 and Δ141-144 on D1^+11^); again, the formation of filaments was not affected (**Figure 3C**). However, filaments in Δ141-144 and Δ291-293 were significantly shorter compared to wild type, which could be due to lack of optimal packing in the D3^0^-D1^+11^ interface and a resulting loss of structural integrity of the filament.

Regardless of how the above-described mutations affected motility, all resulted in the formation of flagellar filaments, suggesting that each FliC mutant was expressed, exported and properly folded by the bacterium. In order to validate that these mutant FliC proteins were similar to wild type, we performed additional *in silico* and biochemical analyses. First, we confirmed that AlphaFold [32, 33] was able to properly predict our crystal structure of PAO1 D2/D3, which it did with an RMSD of 0.573 Å, suggesting that AlphaFold can predict PAO1-like flagellins with high confidence. Then, we used AlphaFold to predict structures of each mutated FliC that we employed in the complementation studies above; each mutant protein was highly structurally similar to our D2/D3 crystal structure (**Figure S4)**. We recombinantly expressed and purified all of the full-length (i.e., inclusive of all domains D0 through D3) FliC deletion mutants, and tested them for proper folding and thermal stability (**Figure S5**). As assessed by size exclusion chromatography, all of them were soluble and monomeric, except for Δ204-205, which was predominantly dimeric. In addition, we subjected each of these deletion mutants to differential scanning fluorimetry and determined that their melting temperatures were all similar to that of the wild type. These data indicate that the changes in swimming motility related to mutations or truncations in the complemented *fliC* genes were not a result of substantial FliC protein structural differences, aggregation or instability.

### Type B FliC provides an advantage in swimming through a viscous environment

Different strains of *P. aeruginosa* contain FliC of either type A or type B and their structures differ significantly. We used the available crystal structure of the type A FliC from the *P. aeruginosa* PAK strain to model L- and R-handed PAK filaments using the helical parameters of the PAO1 filament. Our model suggests that the outer domains of the PAK filament in each protofilament are farther apart than those in PAO1 and do not engage in direct subunit-to-subunit interactions (**Figure 4A**). In order to investigate the consequences of these different filament architectures between the PAO1 and PAK strains, we first compared the motility of PAO1 and PAK wild type strains in liquid and in our agar-based swimming motility assay. Our results showed that while the swimming speeds of the PAK and PAO1 strains in liquid are comparable, the ability of the PAK strain to swim in the semi-solid agar was significantly lower than that of PAO1, with a motile spread of only 20% relative to PAO1, even though both strains have fully formed flagella (**Figures 4B and S2U**).

**Figure 4.**
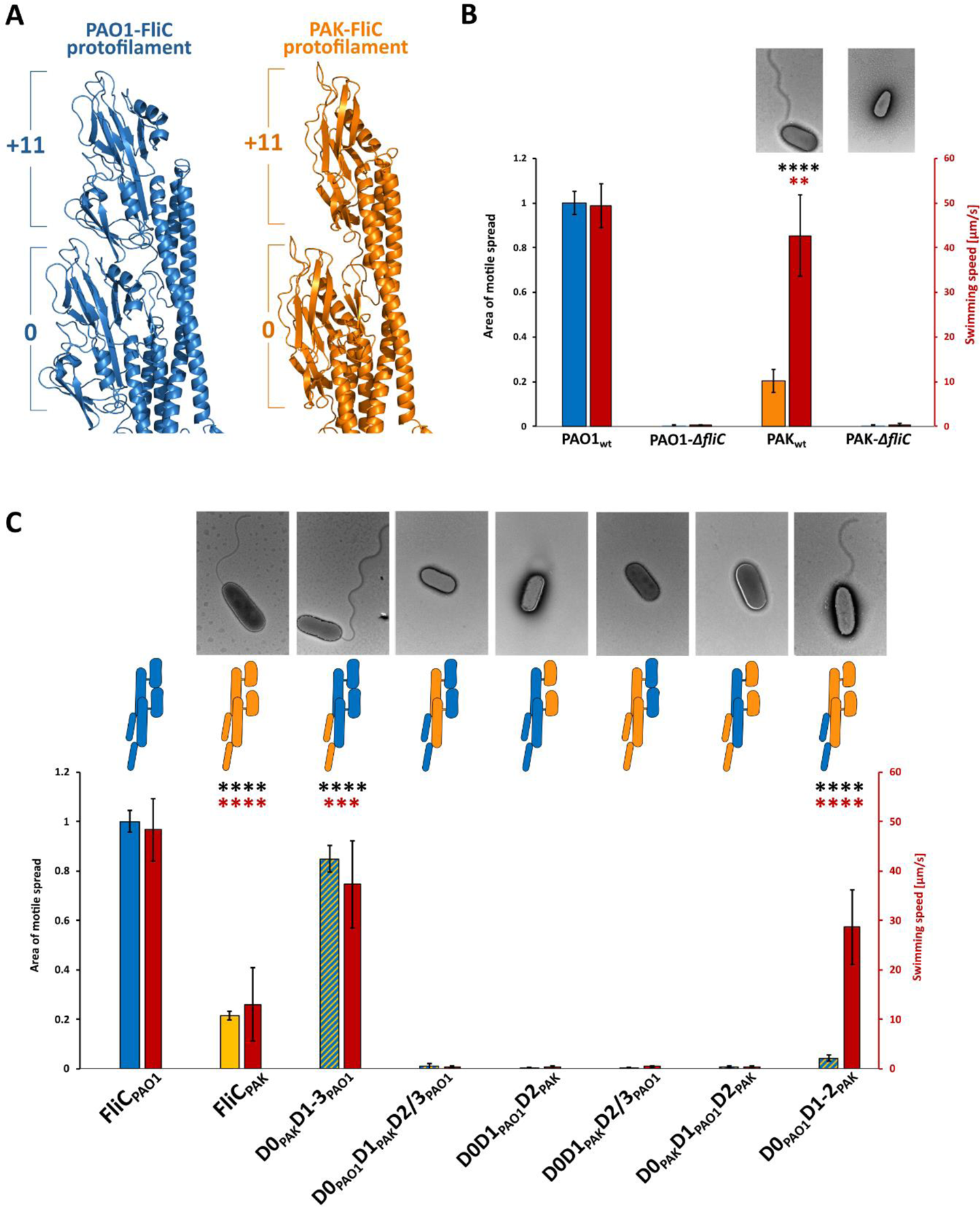
Effects of FliC domain swaps between *P. aeruginosa* PAK and PAO1 strains on swimming motility and filament formation. **A** – Left-handed protofilaments of PAK (orange) and PAO1 (blue) strains superposed. Outer domains in PAK are farther apart and do not engage in contacts; **B** – Swimming motility and negative-stain EM images of the wild type PAO1 and PAK strains; **C** – Swimming motility and negative-stain EM images of PAO1-Δ*fliC* strain complemented with PAK-FliC or FliC in which one or two domains are replaced with equivalent domains of PAK strain. Blue bar (PAO1), yellow bar (PAK) and blue-yellow bars (PAO1-PAK chimeras) represent the motile spread on semi-solid agar. Red bars represent average swimming speed in liquid medium (right vertical axis). Area of motile spread for each strain is normalized to that of the PAO1 wild type strain (**A**) or full-length wild type complemented strain (FliC_WT_) (**B**). (** p < 0.01; *** p < 0.001; **** p < 0.0001).

We next sought to determine whether it was possible to replace type B FliC in PAO1 with type A FliC from PAK. We complemented the PAO1-*ΔfliC* strain with a plasmid-borne type A FliC and tested for motility using the agar-based swimming assay and the presence of filaments by electron microscopy. Our results showed that the PAO1strain complemented by the type B FliC of PAK, like wild type PAK bacteria, had a motile spread of only 20% compared to the strain complemented by type B FliC of PAO1 and formed substantially shorter flagella. It is known that species compatibility between FliC and FliD is required for the filament to be formed [19]. Therefore, we wanted to determine the effect of the replacement of the individual domains in PAO1-FliC with the corresponding domains of PAK-FliC. We generated six constructs in which either one or two domains originated from PAK-FliC and the remainder from PAO1-FliC: D0_PAK_D1-3_PAO1_, D0_PAO1_D1_PAK_D2/D3_PAO1_, D0D1_PAO1_D2_PAK_, D0D1_PAK_D2/3_PAO1_, D0_PAK_D1_PAO_1D2_PAK_, D1_PAO1_D2D3_PAK_. Of these six constructs, only the two strains complemented with *fliC* in which both D1 and the outer domains were from the same strain – D0_PAO1_D1D2_PAK_ and D0_PAK_D1-3_PAO1_ – formed filaments (**Figure 4C**). Replacing the D0 domain of PAO1 with D0 of PAK (D0_PAK_D1-3_PAO1_) led to a decrease in spread of only 15% compared to the knockout strains complemented with wild type *fliC*, comparable to the observed decrease in swimming speed; that is, this complementation resulted in nearly fully restored swimming motility relative to wild type PAO1 (**Figures 4C and S2R-T**). Conversely, replacing D1 and D2/D3 domains of PAO1-FliC with D1 and D2 of PAK (D0_PAO1_D1D2_PAK_) resulted in nearly complete abolition of swimming motility, as indicated by a decrease of motile spread of 95% compared to the wild type. This was in sharp contrast to the 40% decrease of swimming speed of D0_PAO1_D1D2_PAK_ in liquid.

These results indicate that the structure of the type B outer domains and their interactions along the protofilament confer a significant motility advantage to *P. aeruginosa* PAO1 strain in semi-solid agar relative to liquid. Only when the entire network of inter-domain interactions connecting the outer domains to the inner core are present and completely connected in a *P. aeruginosa* PAO1 background is the bacterium competent for swimming through the viscous environment of semi-solid agar.

### Only the C-terminus of *P. aeruginosa* PAO1 FliC can be replaced by the equivalent region of *S.* Typhimurium FliC

As previously mentioned, the FliC outer domains in *Salmonella* adopt a different structure than FliC in *P. aeruginosa* PAO1 and they do not form contacts with neighboring subunits. Conversely, the FliC inner domains, D0 and D1, exhibit 54% identity and 75% similarity. Therefore, we further expanded our functional screen by generating constructs of FliC chimeras optimized for expression in *P. aeruginosa* PAO1 in which we replaced domains D0, D1 and D2/D3 of PAO1-FliC by equivalent domains from *S*. Typhimurium. After complementation of PAO1-Δ*fliC* with these constructs, we observed neither filament formation nor swimming motility (**Figures 5A and S2V-W**). Thus, none of the constructs, including wild type *Salmonella* FliC, could complement for the lack of endogenous FliC in PAO1-Δ*fliC* strain. We next replaced individual helices in D0 or D1, as well as the β-hairpin in D1. Of the resulting six chimeras, only the construct in which the C-terminal helix, Helix 5, of PAO1 was replaced by the *Salmonella* sequence exhibited filament formation (**Figure 5A**). Moreover, its swimming motility was close to that of wild type.

**Figure 5.**
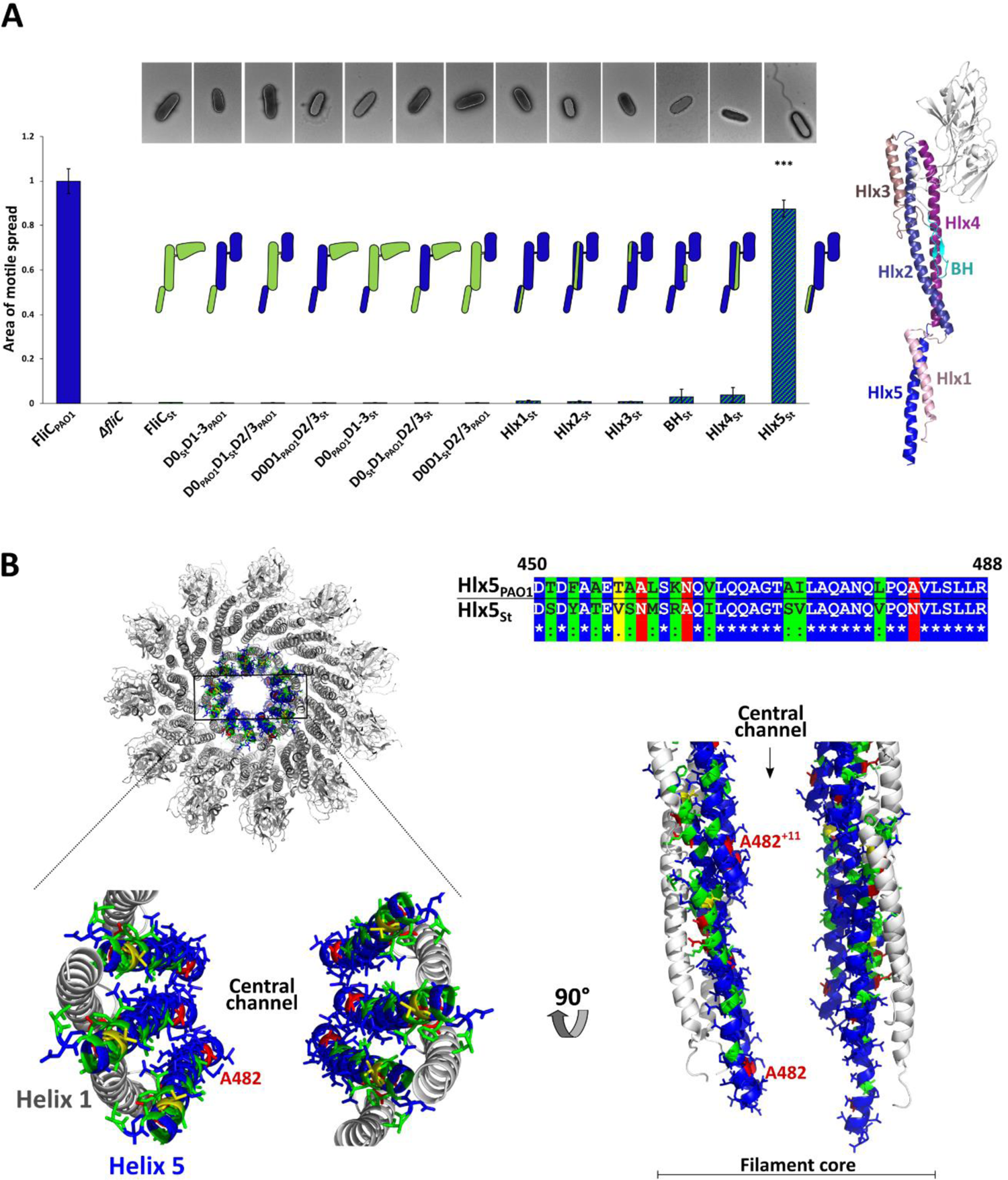
Effects of FliC domain swaps between FliC of *P. aeruginosa* PAO1 and *S*. Typhimurium on swimming motility. **A** – *Left* – Swimming motility analysis and negative-stain EM images of PAO1-*ΔfliC* strain complemented with FliC in which PAO1 domains (blue) or secondary structural elements are replaced with equivalent regions of *S*. Typhimurium (green). *Right* – Secondary structural elements replaced presented on the structure of PAO1-FliC. Hlx – helix, BH – beta-hairpin. Area of motile spread for each strain is normalized to that of the full-length wild type complemented strain(FliC_WT_). (*** p < 0.001); **B** – Cross-section of PAO1 filament showing position of helix 5 in the filament core and sequence alignment of helix 5 from PAO1 and *S*. Typhimurium. Blue - residues that are identical between PAO1 and *S*. Typhimurium. Green – conservative substitutions; yellow and red – non-conservative substitutions.

The C-terminal helix forms part of the inner core of the filament, lining the central channel, making it the only structural part of the filament in contact with the incoming FliC molecule that is being transported to the filament tip (**Figure 5B**). Of the 40 residues that comprise this helix, 14 differ between PAO1 and *S.* Typhimurium, four of which are non-conservative substitutions. Only the non-conserved, partially buried A482 residue (N497 in *Salmonella*) is exposed to the channel lumen. The other three non-conservative substitutions, as well as the majority of conservative substitutions are located on the side of Helix 5 forming contacts with other inner core helices. This suggests that for transport of FliC through the channel, general physico-chemical properties of the channel rather than a specific sequence are important.

### The ridged flagellar filaments characteristic of *P. aeruginosa* PAO1 are found in diverse bacteria

The structure of the flagellar filament of *P. aeruginosa* PAO1 is markedly different from that of *S.* Typhimurium. In *Salmonella*, outer domains D2 and D3 are clearly separated from each other and from neighboring outer domains and do not interact [21]. Conversely, the recently published structure of the *C. jejuni* filament shows a flagellin with three distinct outer domains, with domains D2 and D3 having similar topology to that of D2 and D3 of PAO1 (**Figure 6A**) [25]. The outer domains of *C. jejuni* filament also engage in interactions through D2^0^-D3^+11^ interfaces forming the ridged filament that we observe in PAO1. Recently, AlphaFold structure predictions for over 200 million proteins became available [34]. As mentioned above, AlphaFold prediction of the PAO1 FliC is in excellent agreement with our experimentally obtained crystal structure. Thus, in order to explore the variety of outer domain folds and identify other flagellins with a fold similar to PAO1-FliC, we turned to the newly available database of Alphafold-predicted structures available through Uniprot. We searched for flagellin sequences in Uniprot and visually inspected the Alphafold-predicted structures of those with a size ranging from 450 to 550 amino acid residues.

**Figure 6.**
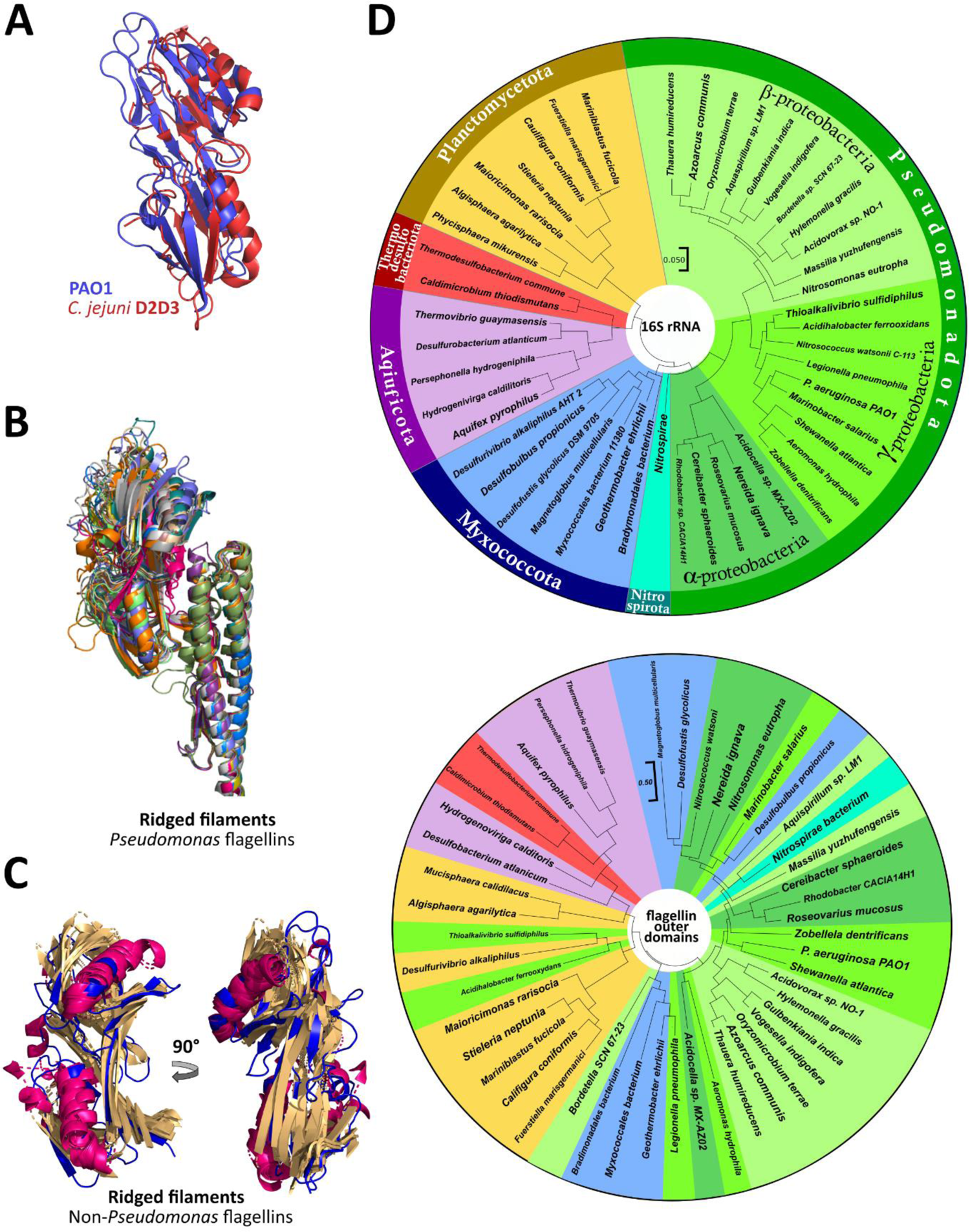
PAO1-like flagellin fold predicted by AlphaFold in different bacterial species. **A** - Superposition of D2 and D3 domains of *C. jejuni* flagellin FlaA and PAO1-FliC; **B** - Superposition of FliC molecules of *Pseudomonas* species other than *P. aeruginosa*; **C** – Superpositions of flagellin outer domains of non-*Pseudomonas* species. Unstructured loop regions were removed for clarity; **D** – Phylogenetic trees based on 16S rRNA (top) and flagellin outer-domain protein sequences (bottom). Species are grouped into bacterial phyla according to the currently accepted ICNP nomenclature.

We first searched for flagellin structures from *Pseudomonas* species other than *P. aeruginosa* and found that at least 10 of these have an outer domain that corresponds to type B FliC (**Figure 6B and S6)**. Next, we expanded our search for flagellin sequences from the entire bacterial kingdom, inclusive of all 42 validated bacterial phyla [35]. Among these bacterial phyla, we found flagellin sequences for 31 of them, the remaining 11 phyla being presumed non-flagellated bacteria. Of the flagellated bacteria, we identified at least 46 species in 6 phyla in which flagellins have a PAO1-like outer domains, including Aquificota, Myxococcotta, Pseudomonadota, Planctomycetota and Thermodesulfobacteriota **(Figures 6C and 6D**). Phylum Pseudomonadota (Proteobacteria) had the largest number of PAO1-like flagellins, with species belonging to the Alpha-, Beta-, and Gamma-proteobacteria classes. We only identified these PAO-FliC-like proteins by visual inspection of Alphafold-predicted structures of FliC proteins of similar length, as these outer domains do not exhibit any significant similarity on the amino acid sequence level (**Figure S6**). These results show that PAO1-like flagellins that form ridged filament are widespread among bacteria. That flagellins of genetically distant bacteria evolved to adopt the same fold of their outer domains despite lacking any measurable sequence similarity suggests that the ridged filament architecture confers advantages to the bacteria that express them, such as for swimming in viscous environments.

## DISCUSSION

Bacterial flagella are highly complex and dynamic molecular machines that translate energy generated by a proton motive force into rotation and, ultimately, thrust in order to propel bacteria through liquid environments. The great majority of each flagellum is composed of many copies of the flagellin protein, in most bacteria called FliC, that form the flagellar filament. The function of the filament is manyfold: primarily, through conformational hand-switching of the FliC subunits it generates thrust by way of an Archimedean screw mechanism; secondarily, it mediates adhesion to host surfaces and other bacteria to promote colonization and biofilm formation, respectively. Flagellar filaments can be neatly divided structurally into the two inner domains, D0 and D1, which form the inner core of the filament, and a diverse number of outer domains, that decorate the exterior of inner core of the filament.

The role of the inner domains is increasingly well understood. They create the flagellar type 3 secretion system (T3SS)-like pore through which flagellin subunits traverse on their way to encountering the FliD filament cap, folding and extending the flagellar filament from its distal end. They are also required for the L- and R-hand-switching that generates thrust in what would otherwise simply be a spinning rod. Since these two functions – growing a filament that subsequently generates thrust – are the minimum requirements for swimming motility, it should come as no surprise that some motile bacteria, such as *B. subtilis*, have evolved to express flagellins containing only D0 and D1 inner domains. Also unsurprising is that these inner domains are highly conserved, in sequence and in structure, throughout flagellated bacteria.

Most flagellated bacteria, however, express flagellins with one or more additional outer domains. Moreover, these outer domains are highly divergent in sequence and, as an increasing number of structural studies have demonstrated, in domain conformation and overall supramolecular architecture. While these remain difficult to fully categorize, some of the observed architectures include splayed, as in *S*. Typhimurium [21, 22], ridged, as in *P. aeruginosa* PAO1 [27], screw-like, as *Sinorhizobium meliloti* [26], and sheathed, as in enterohemorrhagic *E. coli* O157:H7 (EHEC) [26]. While it has been known for decades that flagellin outer domains are involved in adhesion and biofilm formation, their potential role in motility is only recently becoming appreciated. Indeed, a recent study [26] has implicated a role for the dimeric interface between distal D4 domains of the EHEC filament in motility, suggesting that bacteria that inhabit viscous environments are conferred a motility advantage on account of their interconnected outer domains.

Here we show that FliC outer domain interactions: (1) can involve more extensive networked interactions than just intermolecular contacts in outer domain interfaces; (2) provide a motility advantage to bacteria in viscous versus fluid environments; and (3) are more widespread throughout the bacteria kingdom than previously appreciated. Using *P. aeruginosa* PAO1 as a model system of ridged flagellar filaments, we dissected interactions involving its D2 and D3 outer domains and their effects on motility. As a consequence of the architecture of these ridged filaments, these outer domains create a network of interfaces along the protofilament from the inner core, through the outer domains and back to the inner core. By measuring motility of *P. aeruginosa* PAO1 mutant filaments, we found that each one these interfaces – between D1^0^ and D2^0^, connecting the inner core to the outer domains; between D3^0^ and D2^+11^, bridging the outer domains along the protofilament; and between D3^0^ and D2^+11^, reconnecting the outer domains to the inner core – is required for motility. Together, these interfaces create a network of interactions upon which motility depends. Breaking this network at any point results in immotile bacteria.

To demonstrate that FliC outer domains conferred a motility advantage to bacteria in viscous environments, we complemented *P. aeruginosa* PAO1 with PAO1/PAK chimeric filaments. Unlike the ridged filaments in *P. aeruginosa* PAO1, *P. aeruginosa* PAK exhibits splayed filaments in which the outer domains do not make contacts with one another along the protofilament. When we measured motility in liquid, wild type PAO1 and PAK, as well as all chimeras that produced flagella, were similarly motile. Conversely, when challenged with the relatively more viscous soft agar environment, wild type PAO1 was substantially more motile than wild type PAK. Of those chimeras that produced flagella, only that which included the D1, D2 and D3 domains from PAO1 – thereby reconstituting the entire network of interactions between the inner core and outer domains in wild type PAO1 – was similarly motile to wild type PAO1.

We found that ridged flagellar filaments, and consequently the networked interactions between the inner core and outer domains, are not unique to *P. aeruginosa* PAO1. Our computational predictions show that not only are ridged filaments found extensively throughout *Pseudomonas* species, at least 46 species outside of the *Pseudomonas* genus have similarly folded outer domains. Despite similar structures, sequence alignment reveals no significant identity between these outer domains. Thus, while diversity of the outer domain sequences could have resulted in a large number of different outer domain structures, our results suggest that convergent evolution has yielded many paths to a single filament architecture. Due to the laborious method by which we identified these similarly-folded FliC proteins, which relied on visual inspection of individual Alphafold-predicted structures, this undoubtedly represents an undercount; ridged flagellar filaments may be even more widespread throughout bacteria than we have found here. It is also reasonable to believe that the total number of different outer domain structures is smaller than suggested by sequence diversity of outer domains and that filaments could be classified into just a handful of architectures, minimally including: naked (e.g., *B. subtilis*), splayed (e.g., *S.* Typhimurium), ridged (e.g., *P. aeruginosa* PAO1), screw-like (e.g., *S. meliloti*) and sheathed (e.g., EHEC) filaments. Different architectures could reflect the specific needs of distinct groups of bacterial species.

The explanation for the existence of ridged filaments in PAO1 and *C. jejuni*, and consequently in other species, could be in the functional properties of the flagellum. The flagellar motor causes rotation of the filament, creating thrust and enabling bacteria to swim forward. The motor consists of two distinct molecular assemblies, the membrane-embedded rotor, and a torque-generating stator. Stators are composed of a variable number of stator units, with each unit being a complex of the proteins MotA and MotB. In *Salmonella*, the stator consists of 11 subunits. Conversely, the *C. jejuni* motor has up to 17 stator units [36]. This translates to a much higher swimming speed close to 100 µm/s, compared to 25 µm/s of *Salmonella*. *C. jejuni* also swims faster in moderately viscous fluid than in low viscosities [37]. While not much is known about the number of stator units in *P. aeruginosa*, recent cryo-electron tomography analysis showed that its motor is wider than the motor of *S*. Typhimurium and includes additional prominent densities adjacent to the P- and L-rings made of the MotY homolog [38], a protein that is a part of the *Vibrio alginolyticus* Na^+^-driven motor. Additionally, *P. aeruginosa* has two types of stator units, MotA/MotB for swimming in low viscosity, and MotC/MotD for swimming in higher viscosity [39]. All these specificities of the *P. aeruginosa* motor would result in generation of greater maximum torque, allowing bacteria to swim at higher velocities through more viscous media [38].

Although the maximum swimming speed of *P. aeruginosa* of 50 µm/s is half the speed of *C. jejuni*, it is still twice that of *S*. Typhimurium [40]. Both *P. aeruginosa* and *C. jejuni* have polar flagella. Due to the higher force and load to which their filament is subjected, it is reasonable to expect a much tighter packing to preserve its structural integrity. As such, outer domains with extensive inter-domain interactions could stabilize the filament while swimming in the viscous environment, while the lack of connectedness of the PAK outer domains together with its poor relative motile spread in viscous environments suggests that its filament architecture is poorly adapted to viscosity. Unlike monotrichously flagellated *Pseudomonas* species, *S*. Typhimurium is peritrichously flagellated and its individual filaments bundle during straight forward swimming. A looser outer domain structure could serve as a mechanism for easier bundling and unbundling when the change of direction is needed.

## MATERIALS AND METHODS

### D2-D3 purification and crystallization

The sequence of *fliC* from *P. aeruginosa* PAO1 between residues 178 and 395 containing D2 and D3 domains (FliC_D2D3_) was cloned into a pGEX-5x-2 plasmid with the GST tag on N-terminus followed by a TEV protease recognition site. The plasmid was introduced into BL21(DE3) cells. For the native protein expression, cells were grown in LB medium and the expression was induced with 1 mM IPTG, after which the cells were grown for another 3 hours at 37 °C. Selenomethionine-labeled (SeMet) FliC_D2D3_ was expressed for 6 hours after the induction at 37 °C in M9 medium containing the amino acids lysine (100 mg/L), phenylalanine (100 mg/L), threonine (100 mg/L), isoleucine (50 mg/L) and valine (50 mg/L) to suppress methionine synethsis, and 60 mg/L of selenomethionine. The cells were harvested, resuspended in PBS and sonicated. The soluble fraction of the lysate was applied to the Glutathione Sepharose beads (GE Healthcare), washed and eluted with PBS containing 25 mM of reduced glutathione. GST tag was removed by overnight incubation with TEV protease followed by affinity chromatography using Ni-NTA beads to remove TEV protease, and size-exclusion chromatography (Superdex 200, GE Healthcare). SeMet-FliC_D2D3_ crystallized in 1.9 M ammonium sulfate and 0.1 M sodium Citrate pH 5.5, while native FliC_D2D3_ crystallized in 2 M ammonium sulfate and 0.1 M Tris pH 7.5.

### D2-D3 crystallization and structure determination

Data sets were collected on the BL12-2 beamline of the Stanford Synchrotron Radiation Lightsource (SSRL) equipped with the Pilatus 6M detector (Dectris). For the selenomethionine-labeled FliC, we collected 3 data sets at peak (λ=0.97929 Å), high-energy remote (λ=0.9116 Å) and inflection (λ=0.9795 Å) wavelengths. Data were indexed, integrated and scaled using the XDS program package [41]. The MAD phasing was performed using SHELX program in CCP4i package [42] and the initial model was built with Buccaneer [43]. The native dataset was collected at λ=0.97946 Å. Data were phased by molecular replacement with MOLREP [44], using the previously obtained model from the anomalous scattering. The final model at 1.47 Å resolution was refined in several rounds using REFMAC5 [45] and manual editing in Coot [46], and has been deposited to the PDB with the entry code 8ERM.

### Whole-atom filament reconstitution

The workflow for building the full-length FliC molecule and subsequent whole-atom reconstruction of both L- and R-handed filaments was the same, as follows. Previously obtained electron density map of PAO1 L- and R-handed filaments and the models of the filament core served as a starting point. First, manual rigid-body fit of the D2/D3 crystal structure was performed in UCSF Chimera [47] using a filtered electron density map and then further refined by using Chimera’s Fit in map function. The missing residues connecting D1 and D2/D3 were built using Sali lab’s ModLoop [48], resulting in a full-length FliC monomer Filaments were reconstituted by applying helical symmetry operators in Phenix [31], and refined using Phenix’s Real-space refinement program.

### *ΔfliC*-PAO1 complementation

Wild-type and *ΔfliC* strains *P. aeruginosa* PAO1 were obtained from the Manoil Lab at the University of Washington, while *P. aeruginosa* PAK strains were kindly provided by Joanna Goldberg from Emory University. Different constructs of *fliC* genes were cloned into the pBBR1 plasmid, which was then transformed into the knockout strains by electroporation [49].

### Swimming assays

Swimming motility assays of *Pseudomonas aeruginosa* strains were performed as described in [50]. Plates were incubated at 37 ⁰C for approximately 20 hours. The area of each bacterial swim circle was quantified using the software ImageJ [51]. Ten replicates were performed for each complemented strain, and the average and standard deviations were determined. Statistical significance was determined by Brown-Forsythe and Welch ANOVA test followed by a Dunnett’s T3 multiple comparison test using GraphPad Prism version 9.2.0 for Windows (GraphPad Software, San Diego, California USA, www.graphpad.com).

### EM analysis

Bacteria were grown in Luria Broth liquid culture overnight at 37°C and immobilized on a Formvar grid (Electron Microscopy Sciences) for 1 min 30 sec and the samples were negatively stained with 0.5% (w/v) Phosphotungstic acid (PTA) and imaged using a FEI Talos 120 KV electron microscope.

### Swimming velocity determination

Cells from overnight culture were diluted into the LB medium to an OD600 of 0.05 and incubated for 30 minutes at 37 °C. Cells were observed and recorded in a phase-contrast mode on Lionhear FX automated microscope (Biotek) using a wet mount technique. The recordings were analyzed using the TrackMate plugin [52] in the image processing package Fiji [53]. Average speed for each strain was determined using 30 fastest tracks.

### Protein purification and differential scanning fluorimetry (DSF)

Genes of the wild type FliC and deletion mutants were subcloned into pET28-a vector with C-terminal polyhistidine tag. Plasmids were transformed into *E. coli* BL21(DE3) cells, grown in LB medium at 37⁰ C until OD_600_=0.6. After induction with 1 mM IPTG, cells were grown for another 4 hours at 37⁰ C. Clarified lysate was applied to NiNTA resin (Qiagen) and eluted with 500 mM imidazole in PBS buffer, followed by size-exclusion chromatography using Superdex 200 Increase 10/300 GL column (Cytiva). For DSF analysis, 5 µM of protein in PBS was mixed with SYPRO Orange dye (ThermoFisher) to a final concentration of 25X. The experiment was performed on a real-time thermal cycler for a temperature range between 25⁰ and 90⁰ C.

---

**Supplemental figure S1.**
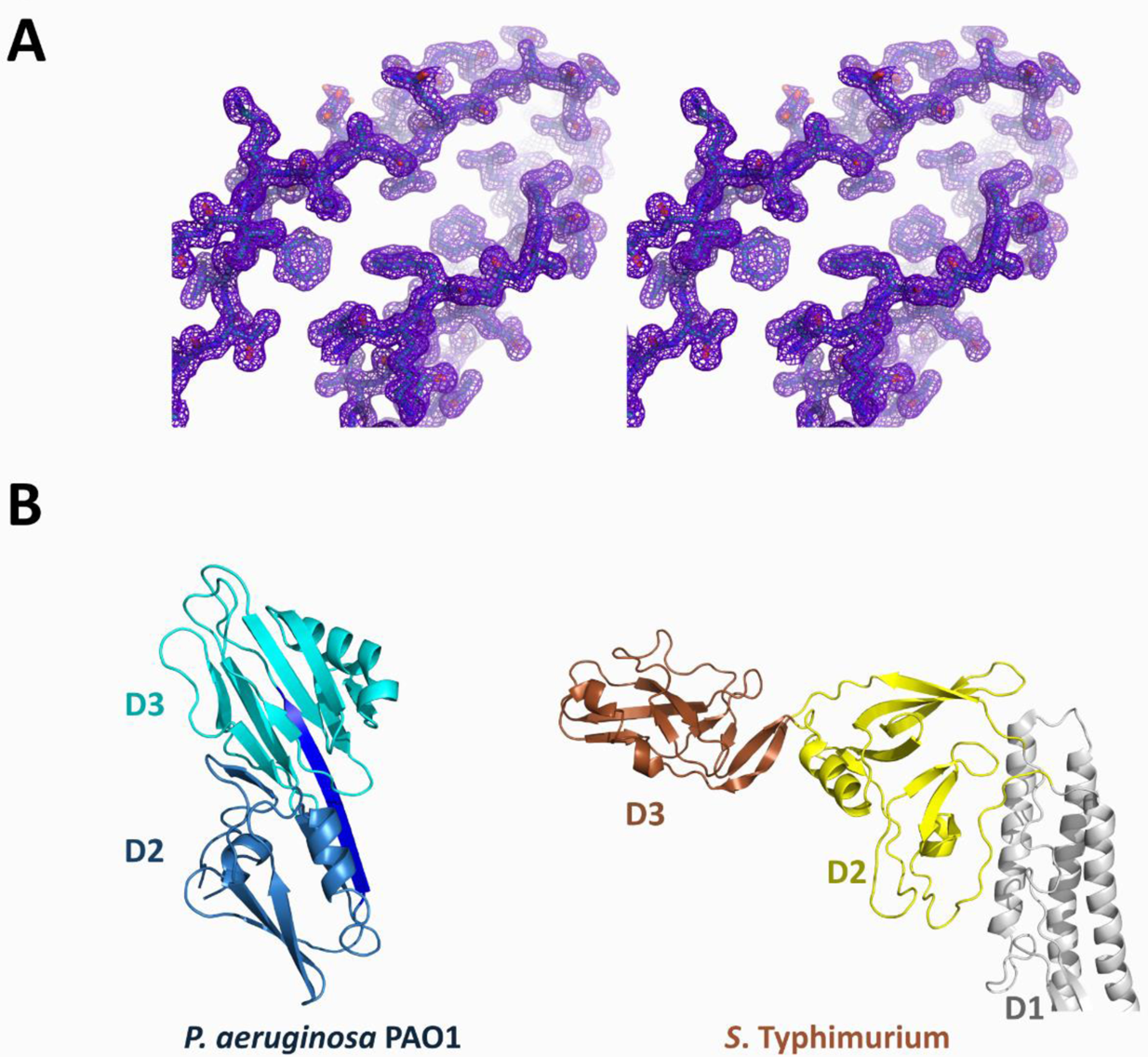
**A** – Stereo image of a portion of the 2Fo - Fc electron density map of native FliD_D2D3_ 2Fo - Fc electron density map. **B** – Comparison of D2 and D3 domains of *P. aeruginosa* PAO1 and *S*. Typhimurium.

**Supplemental figure S2.**
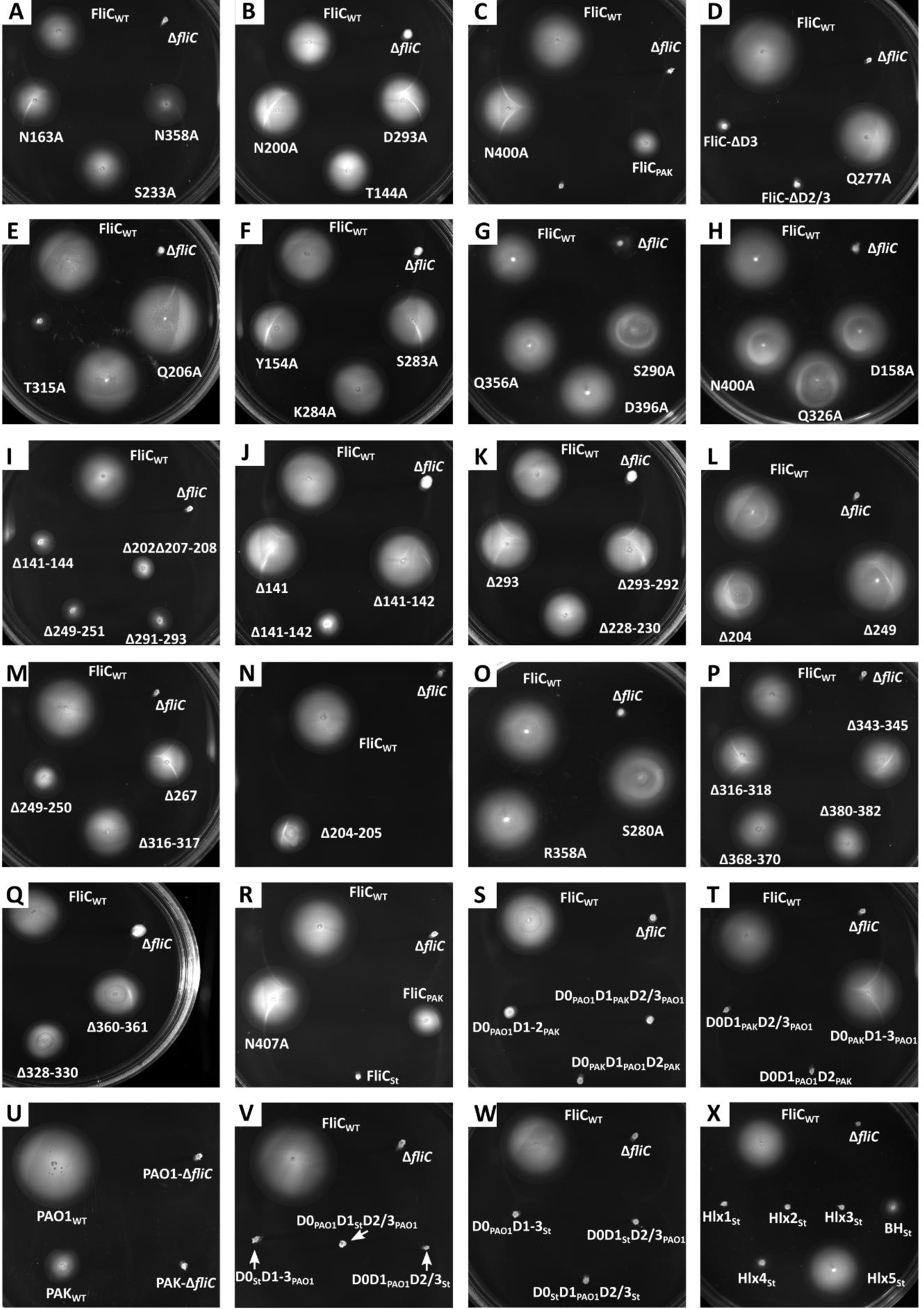
Agar plates showing motile spread of *P. aeruginosa* PAO1-Δ*fliC* strain complemented with different FliC mutants.

**Supplemental figure S3.**
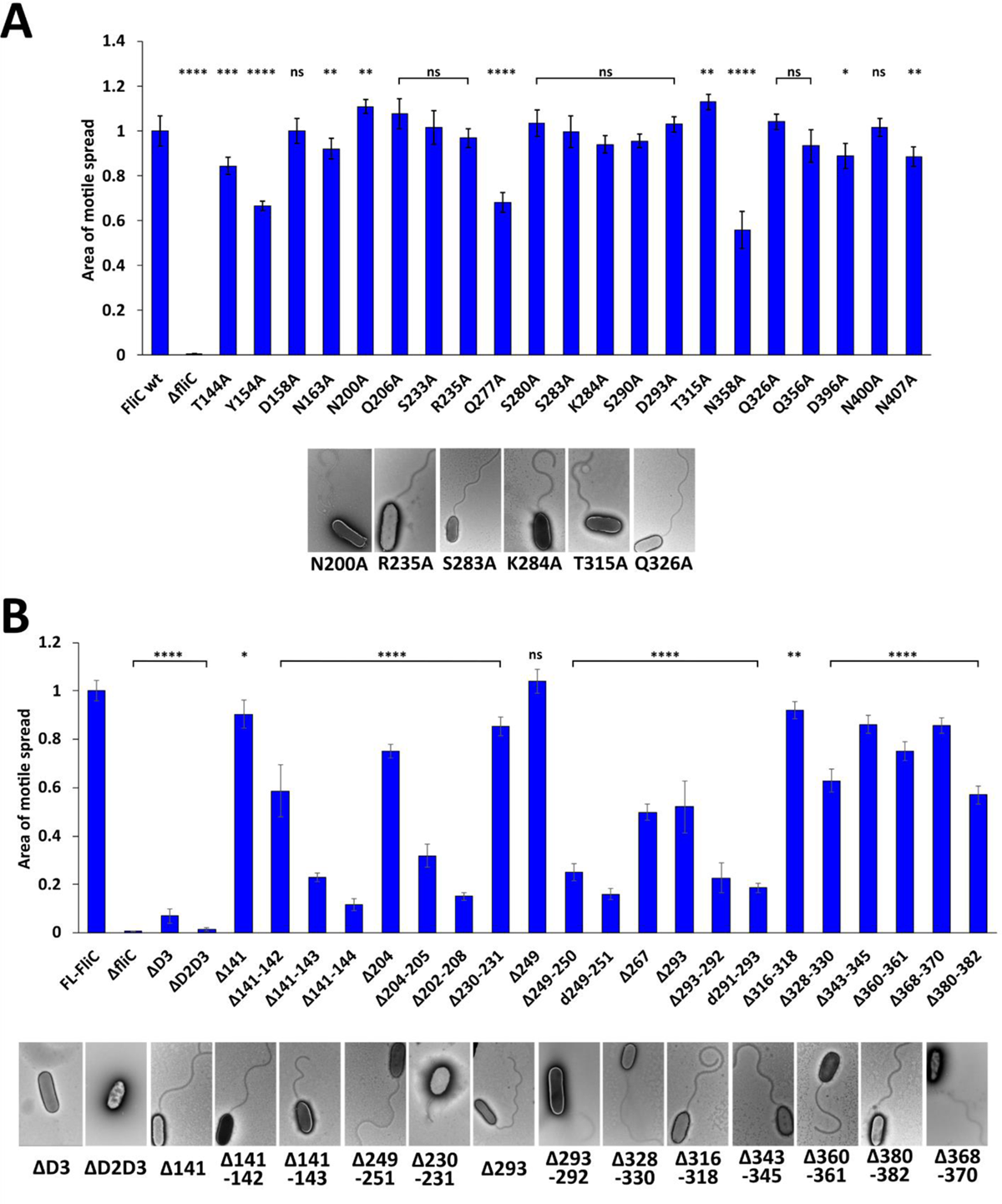
Motile spread of all alanine mutants (**A**) and deletion mutants (**B**) tested. Negative-staining EM images presented are the images of mutants not included in Figure 3.

**Supplemental figure S4.**
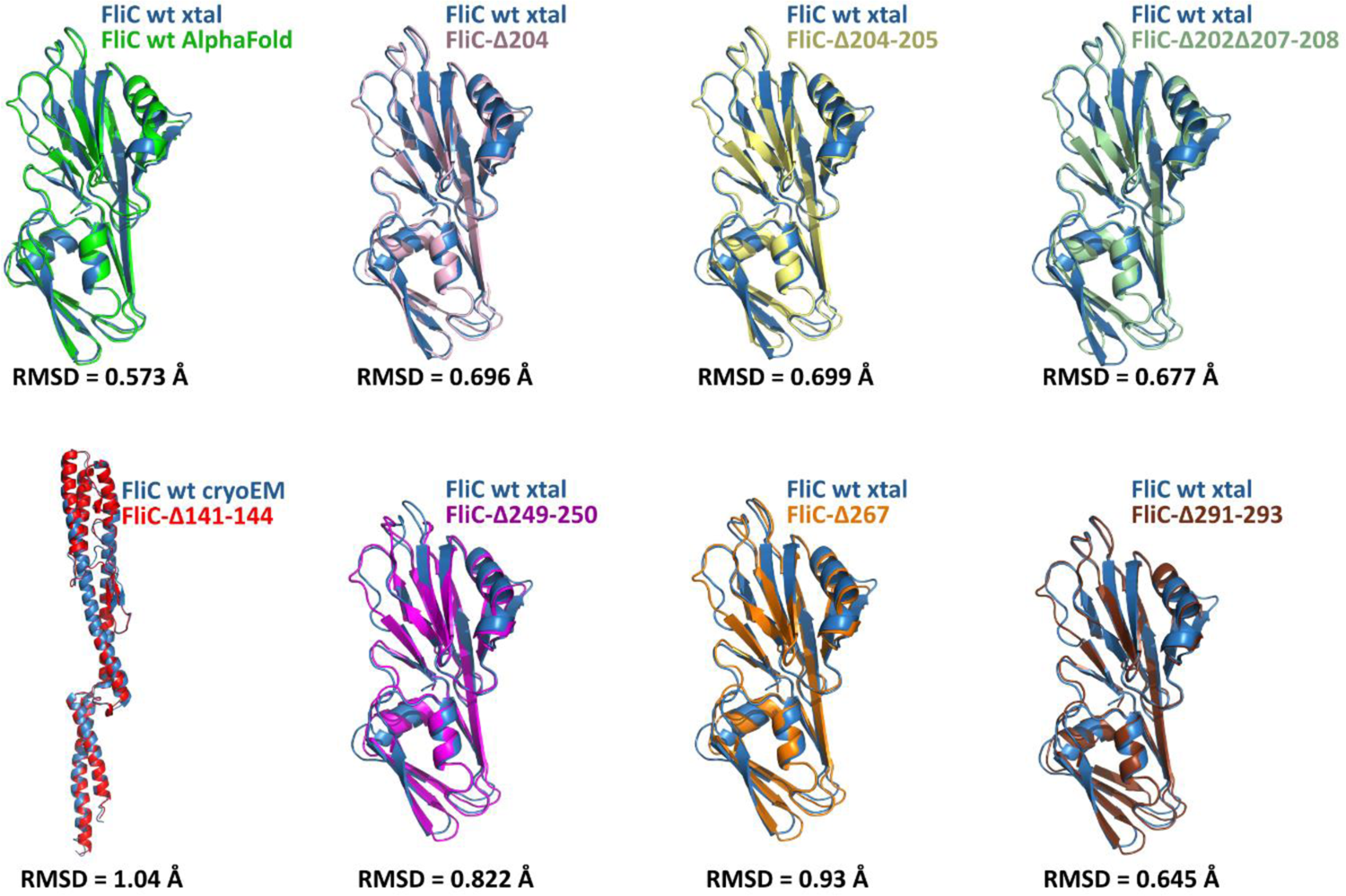
Superposition of PAO1-FliC D2/D3 crystal structure and structures of FliC mutants predicted by AlphaFold.

**Supplemental figure S5.**
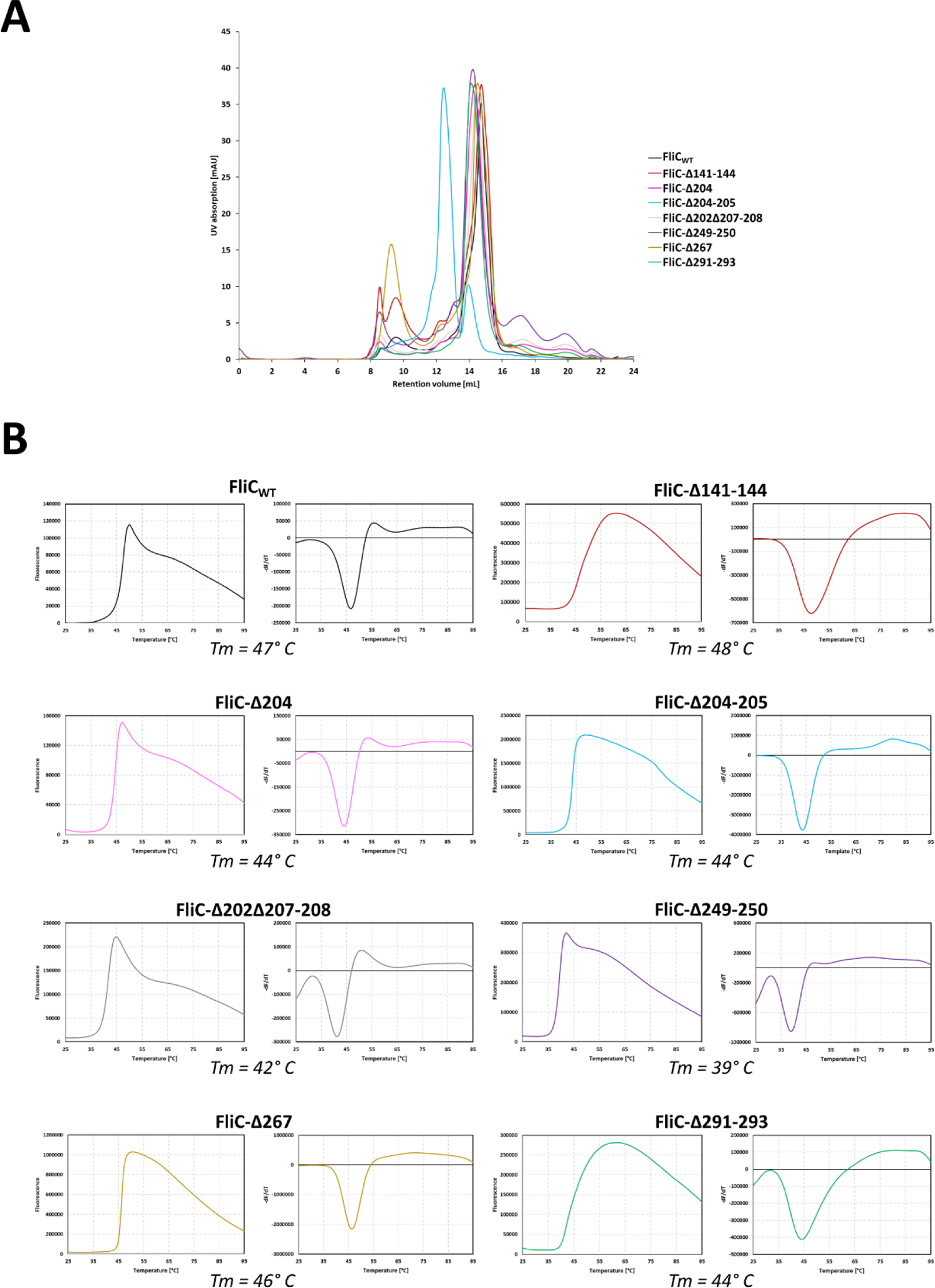
**A** – Size-exclusion chromatogram for recombinantly purified wild type FliC and 7 deletion mutants; **B** – Fluorescence emission and the first derivative obtained from differential scanning fluorimetry (DSF) experiment.

**Supplemental figure S6.**
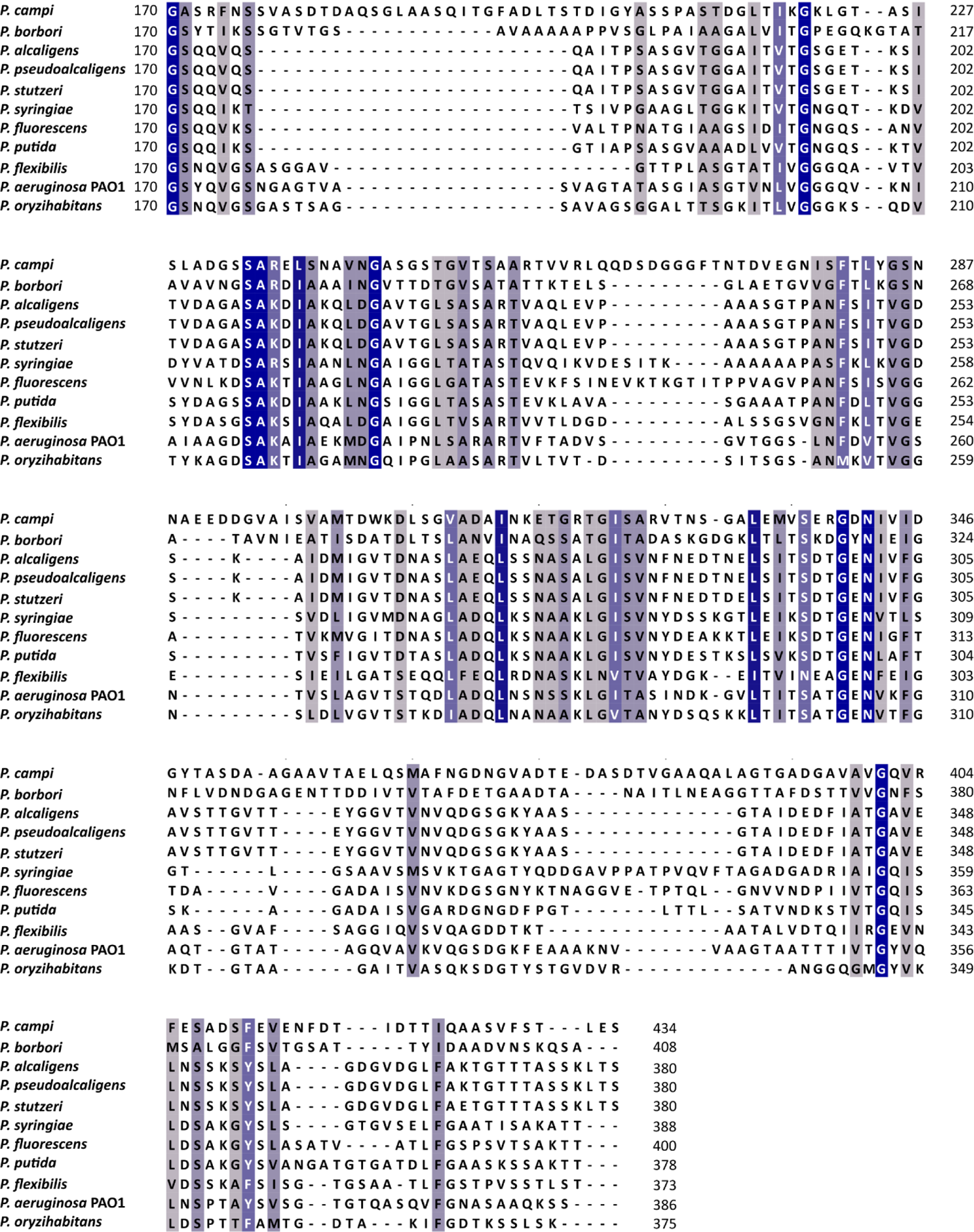
Protein sequence alignment of outer domains of P. aeruginosa PAO1and 10 other species of the genus *Pseudomonas* with PAO1-like flagellin

**Supplemental figure S7.**
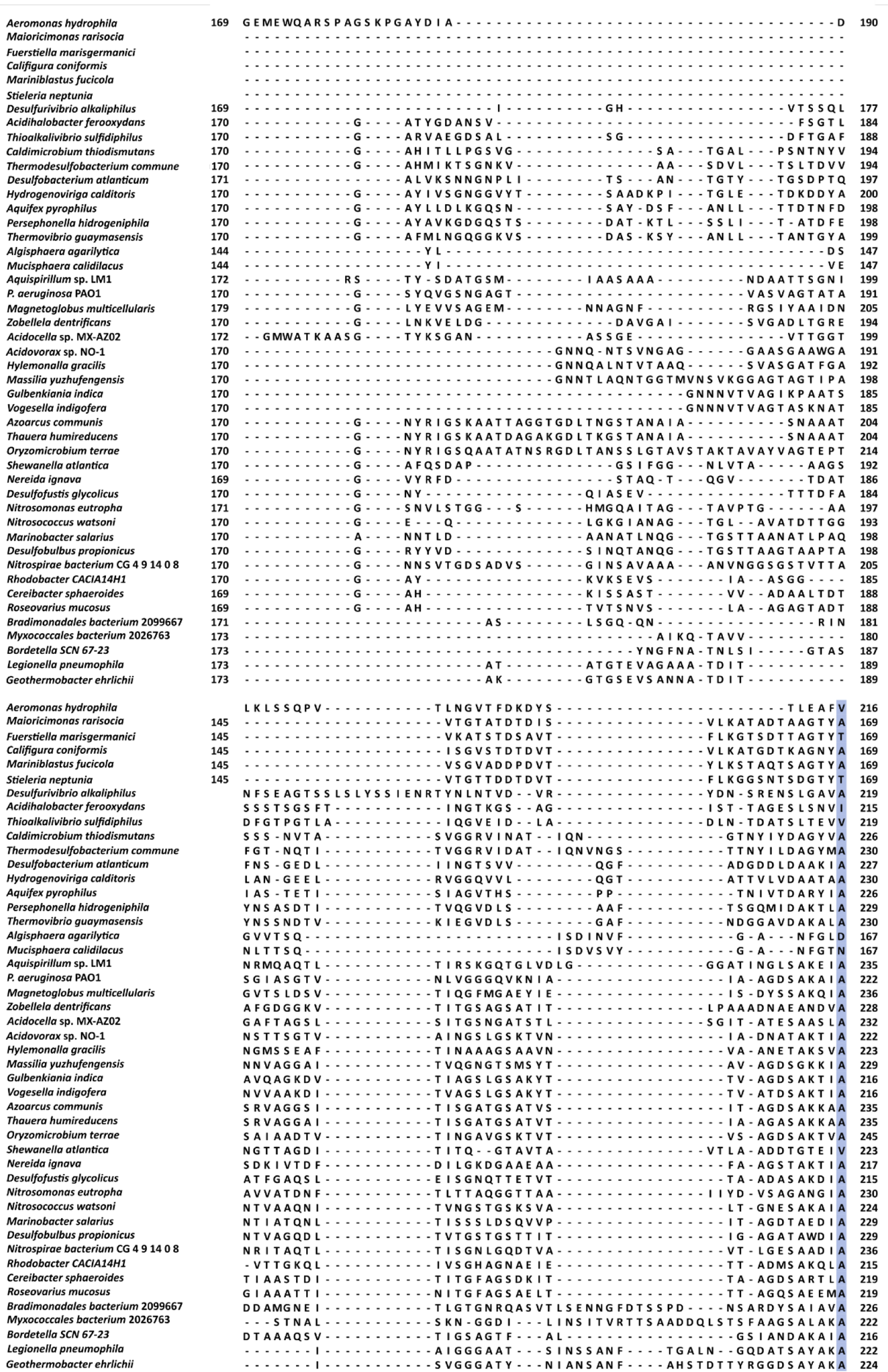
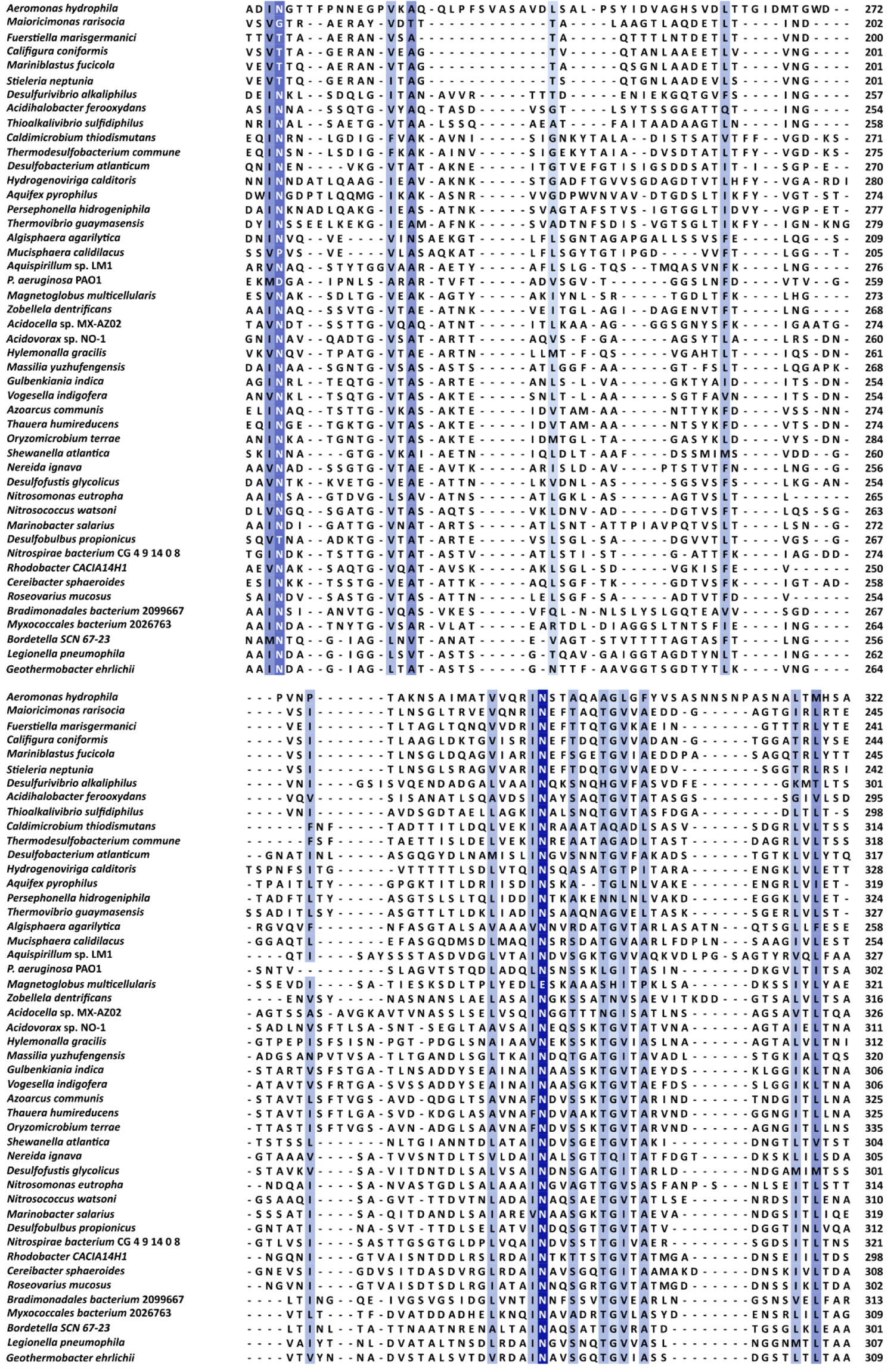
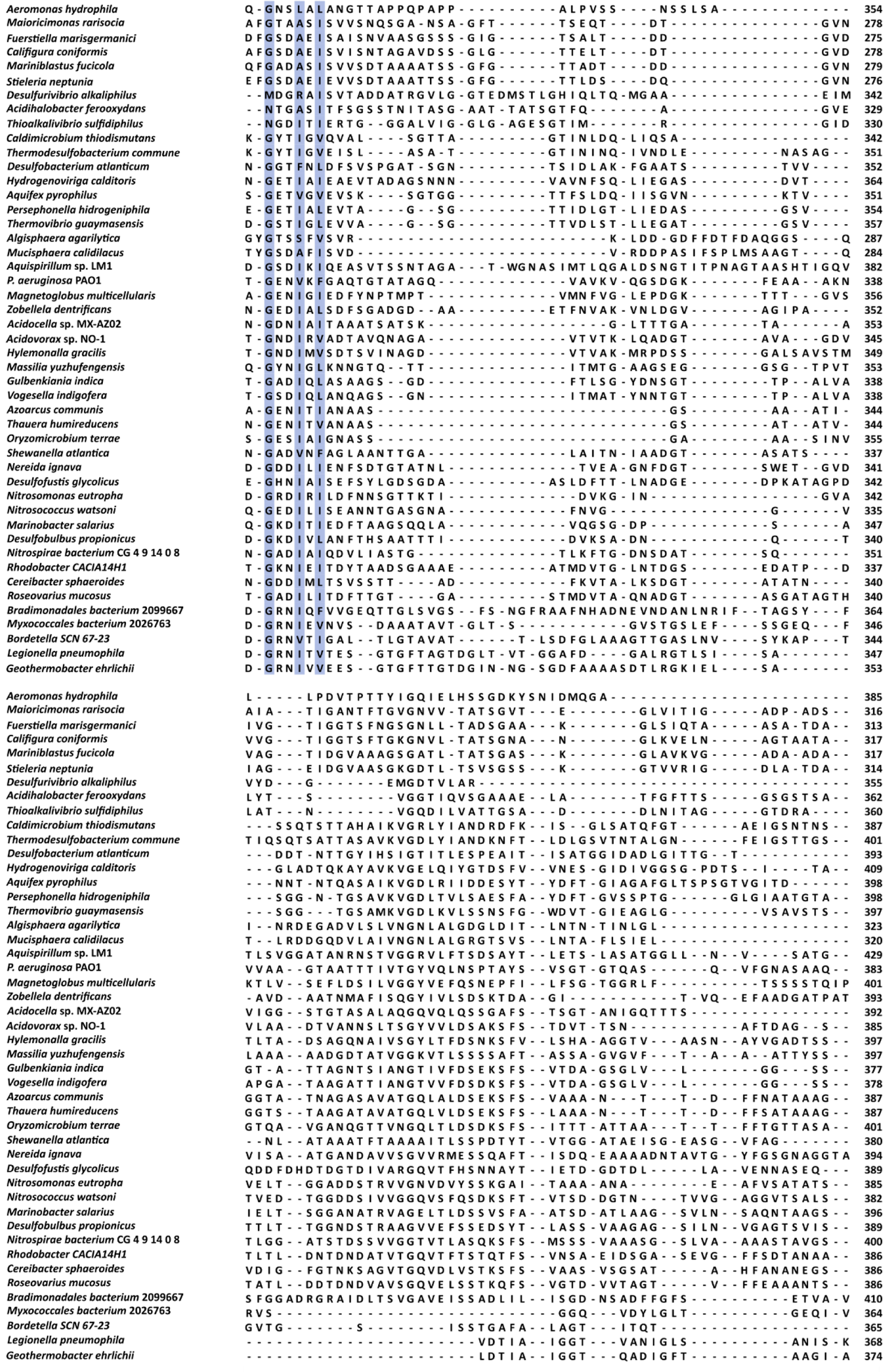
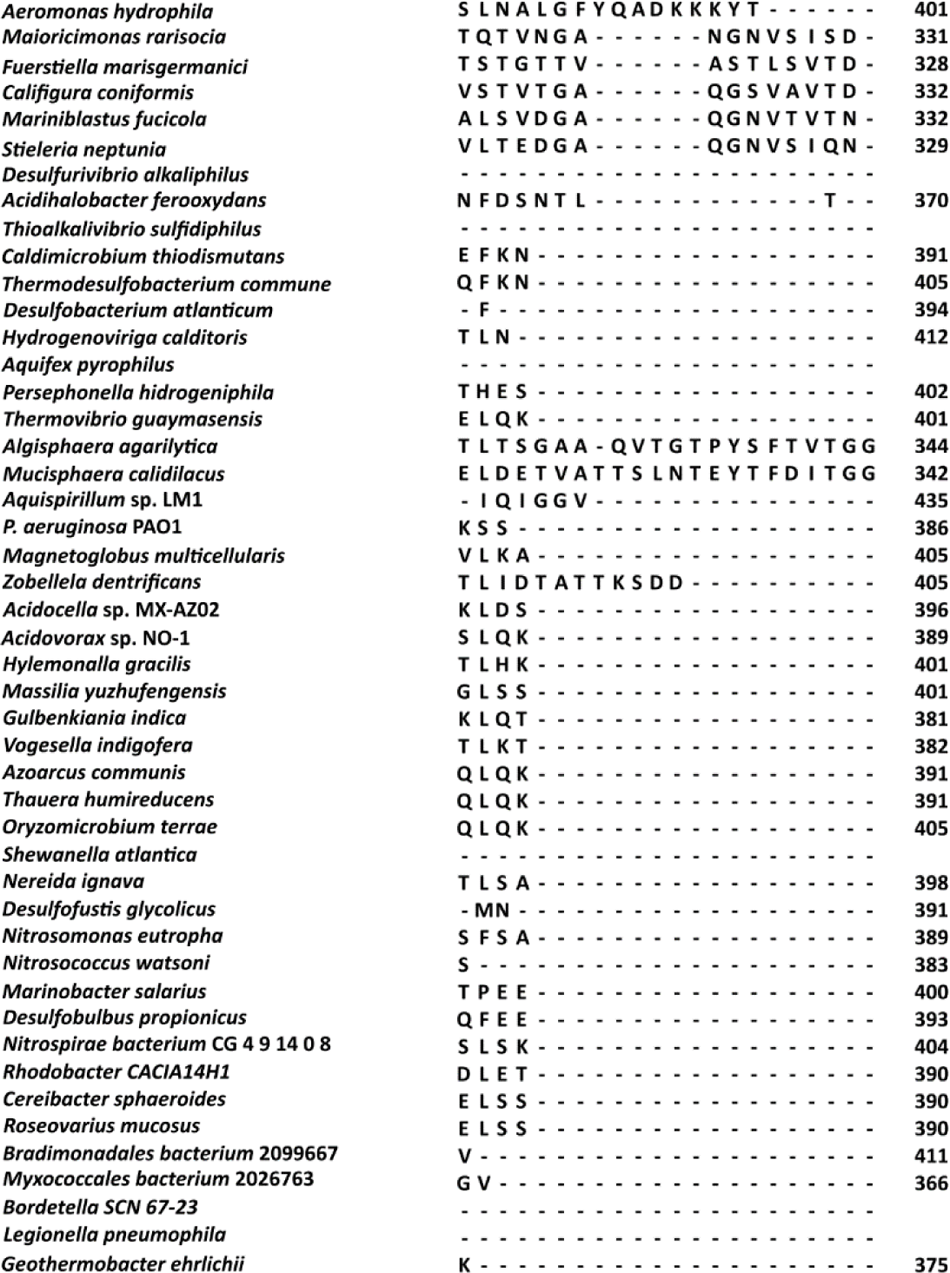
Protein sequence alignment of outer domains of *P. aeruginosa* PAO1 and 46 species with PAO1-like flagellin.

**Supplemental Table S1.** Data collection, phasing and refinement statistics for MAD (SeMet) structures

|  | Native FliC-D2D3 | SeMet FliC-D2D3 |  |  |
| --- | --- | --- | --- | --- |
| **Data collection** |  |  |  |  |
| Space group | P 1 2<sub>1</sub> 1 | P 1 2<sub>1</sub> 1 |  |  |
| Cell dimensions |  |  |  |  |
| *a*, *b*, *c* (Å) | 53.49, 36.49, 87.45 | 53.6, 36.5, 87.7 |  |  |
| α, β, γ (°) | 90.00, 97.30, 90.00 | 90, 97.185, 90 |  |  |
|  |  | *Peak* | *Inflection* | *Remote* |
| Wavelength |  | 0.97929 | 0.97949 | 0.91162 |
| Resolution (Å) | 31.67-1.47 | 36.5 – 1.6 | 36.5 – 2.0 | 36.5 – 2.2 |
| *R*<sub>sym</sub> or *R*<sub>merge</sub> | 0.091 (0.816) | 0.098 (0.510) | 0.075 (0.166) | 0.109 (0.194) |
| *I* / $\sigma I$ | 11.87 (1.52) | 8.9 (2.4) | 14.3 (7.7) | 13.4 (6.3) |
| Completeness (%) | 98.21 (93.99) | 97.7 (95.0) | 90.4 (83.0) | 85.7 (78.0) |
| Redundancy | 6.7 (6.0) | 1.5 (1.6) | 1.6 (1.7) | 1.5 (1.6) |
| **Refinement** |  |  |  |  |
| Resolution (Å) | 86.74-1.47 | 35.95-1.6 |  |  |
| No. reflections | 53148 | 44280 |  |  |
| *R*<sub>work</sub> / *R*<sub>free</sub> | 0.175 / 0.2021 | 0.2425 / 0.2717 |  |  |
| No. atoms |  | 3001 |  |  |
| Protein | 2739 | 2661 |  |  |
| Ligand/ion | 22 |  |  |  |
| Water | 346 | 340 |  |  |
| *B*-factors | 18.53 | 19.06 |  |  |
| Protein | 17.51 | 16.99 |  |  |
| Ligand/ion | 40.17 |  |  |  |
| Water | 25.27 | 35.2 |  |  |
| R.m.s deviations |  |  |  |  |
| Bond lengths (Å) | 0.02 | 0.02 |  |  |
| Bond angles (°) | 1.94 | 2.5 |  |  |
| Ramachandran plot |  |  |  |  |
| Favored (%) | 99.75 |  |  |  |
| Allowed (%) | 0.25 |  |  |  |
| Disallowed (%) | 0.00 |  |  |  |

\*Each structure was determined using data collected from a single crystal. \*Values in parentheses are for highest-resolution shell.

**Supplemental table 2.** Refinement statistics for whole-atom models of *P. aeruginosa* PAO1 filament

|  | Left-handed | Right-handed |
| --- | --- | --- |
| *B* factors (Å<sup>2</sup>) |  |  |
| Protein | 95.66 | 59.27 |
| Ligand |  |  |
| R.m.s. deviations |  |  |
| Bond lengths (Å) | 0.003 | 0.005 |
| Bond angles (°) | 0.601 | 0.819 |
| Validation |  |  |
| MolProbity score | 2.64 | 2.52 |
| Clashscore | 17 | 17.24 |
| Poor rotamers (%) | 5.15 | 2.22 |
| Ramachandran plot |  |  |
| Favored (%) | 94.6 | 91.03 |
| Allowed (%) | 5.40 | 8.76 |
| Disallowed (%) | 0.00 | 0.21 |

